# Single-molecule light-sheet microscopy with local nanopipette delivery

**DOI:** 10.1101/2020.09.25.313973

**Authors:** B. Li, A. Ponjavic, W. H. Chen, L. Hopkins, C. Hughes, Y. Ye, C. Bryant, D. Klenerman

## Abstract

Detection of single molecules in biological systems has rapidly increased in resolution over the past decade. However, delivery of single molecules has remained a challenge. Currently there is no effective method that can both introduce a precise amount of molecules onto or into a single cell at a defined position, and then image the cellular response. Here we have combined light sheet microscopy with local delivery, using a nanopipette, to accurately deliver individual proteins to a defined position. We call this method local delivery selective plane illumination microscopy (ldSPIM). ldSPIM uses a nanopipette and the ionic feedback current at the nanopipette tip to control the position from which molecules are delivered. The number of proteins delivered can be controlled by varying the voltage applied. For single-molecule detection, we implemented single-objective SPIM using a reflective atomic force microscopy cantilever to create a 2µm thin sheet. Using this setup, we demonstrate that ldSPIM can deliver single fluorescently-labeled proteins onto the plasma membrane of HK293 cells or into the cytoplasm. Next, we deposited aggregates of amyloid*-*β, which causes proteotoxicity relevant to Alzheimer’s disease, onto a single macrophage stably expressing a MyDD88-eGFP fusion construct. Whole-cell imaging in 3D mode enables live detection of MyDD88 accumulation and formation of MyDDosome signaling complexes, as a result of aggregate-induced triggering of toll-like receptor 4. Overall, we demonstrate a novel multifunctional imaging system capable of precise delivery of single proteins to a specific location on the cell surface or inside the cytoplasm and high-speed 3D detection at single-molecule resolution within live cells.

**Statement of Significance:** This paper describes and validates a new method to study biological processes based on the controlled local delivery of molecules onto or into the cell, combined with single molecule imaging using light sheet microscopy. we not only demonstrate the instrument’s capability of delivering controlled numbers of molecules to a defined position, down to the level of single molecules, but also its potential in study of the triggering of the innate immune response by protein aggregates, a key process in the development of neurodegenerative diseases such as Alzheimer’s disease. The same approach could be applied to a wide range of other important biological processes allowing them to be followed in live cells in real-time, hence it will be of great interest to the biophysical community.

## Introduction

As research in cell biology is advancing towards ever more complex and detailed studies, a major challenge is to dissect the spatiotemporal-dependent interactions that are critical to cellular functions and this requires the development of new instrumentation to overcome the limitations of current methodologies. For example, the activation of inflammatory pathways upon detection of pathogens and cytokines (1–4) by the innate immune response leads to rapid the formation of a signalling complex at the cell membrane. On immune cells such as macrophages, Toll-like receptors (TLRs) are present to screen for hostile agents(1, 5). Detection of a pathogen triggers dimerization of TLRs, which then recruits proteins, including MyD88, to perform downstream signal transduction(4, 6–11). MyD88 is the main protein in

MyDDosomes, a key supramolecular organising center assembled transiently to activate a cascade of signaling events, most notably inflammatory responses associated with the NF-κB pathway(10, 12, 13). We and others have shown such transient assembly of MyDDosome triggered by lipopolysaccharide (LPS) can be detected on the surface of fixed cells using advanced microscopy, such as total internal reflection fluorescence (TIRF) imaging techniques(14). However, this approach has several disadvantages. TIRF microscopy is limited to image a range ∼200 nm of the water-coverslip interface and the molecules cannot be delivered to the basal surface of the cells that are being imaged. Therefore, innovative approaches to locally and repetitively trigger key biological processes in a controlled way to a single cell, combined with single-molecule imaging to follow the cellular response, are now required to study both TLR4 signaling and receptor signaling in general.

To address the imaging limitation of TIRF, selective plan illumination microscopy (SPIM) has been developed to greatly improve 3D single-molecule imaging in live cells in terms of image acquisition rate and spatial depth. In SPIM, a thin layer within the biological specimen is selectively excited orthogonal to the detection path, minimizing both out-of-focus signals, phototoxicity and photobleaching during imaging(15–18). Several previous SPIM-based studies have demonstrated sensitive detection of receptors on the cell membrane and cell components such as mitochondria in the cytoplasm(19–21). The main difficulty with implementing SPIM is the introduction of the light-sheet into the biological samples and numerous solutions have been developed to minimise this complication(21–25). Single-objective SPIM(20) is notable as it does not prevent sample access, which in our case enables the introduction of a nanopipette delivery system. We have previously developed a nanopipette positioning-and-probing technique – scanning ion conductance microscopy (SICM). Changes in pipette current provide a real-time feedback signal that allows a nanopipette to hover over or penetrate into a biological sample(26). The nanopipette can be used for the controlled voltage or pressure driven delivery of small molecules, proteins and antibodies to defined positions on cell surface or into cytoplasm down to the level of single molecules(27–29).

Here we demonstrate an innovative microscopy design that combines single-molecue light-sheet imaging and nanopipette delivery, which we have named local delivery selective plan illumination microscopy (ldSPIM). This design enables observation of real-time molecular events within the cytoplasm and on the apical surface of live cells initiated by accurate macromolecule delivery using a nanopipette. We demonstrate that our ldSPIM system is capable of accurate delivery of a precise number of proteins onto or into cells. We further use ldSPIM to show that toxic aggregates delivered to the apical surface of macrophages stimulate sustained MyDDosome formation, which we track over time in 3D. Our ldSPIM system should facilitate new single molecule studies of cellular function and dysfuntion.

## Material and method

### Nanopipette fabrication

Nanopipettes were fabricated from quartz capillaries (Sutter Instrument outer diameter 1mm inner diameter 0.5 mm with filament, CA, USA) using a laser pipette puller (P-2000, Sutter Instruments, Novato, CA). Two sets of parameters were used to pull pipettes with two different opening sizes. The size of nanopipettes pulled using these parameters were characterised using scanning electron microscopy (SEM) as previously (26, 30, 31). Detail information of pulling parameters are described in supplementary method and material.

### Wheat germ agglutinin delivery on glass and fixed cell surfaces

Wheat germ agglutinin (WGA) is a positively charged molecule that it is able to binds to glycoconjugates on cell membrane as well as the negatively-charged glass surface so we choose WGA as our model molecule to characterize the delivering function of ldSPIM. When cells are fixed, the glycoconjugates on the cell membrane become immobile so that deposited WGA molecules should be static, making it suitable for single molecule analysis. A nanopipette with ∼100 nm aperture was backfilled with 8 μL positively charged Alexa-555 labelled WGA (Thermo Fisher) at a concentration of 9 μM for bulk delivery and 450 nM for single-molecule delivery in PBS buffer, with an Ag/AgCl electrode. The pipette was immersed into a bath of PBS buffer, which contained an untreated glass surface or cell sample, while another Ag/AgCl electrode served as the reference electrode. A −100mV potential bias from the patch clamp amplifier was applied between the electrodes inside the nanopipette and bath preventing WGA molecules escaping from the nanopipette. This generated a current of approximately 0.4 nA. A custom LabVIEW (National Instrument 2014) program implemented on a Field Programmable Gate Array (FPGA, National Instruments) was used to control the delivering process. The nanopipette was first moved down to a position ∼15 μm above the sample surface using the white light microscope. Then the piezo moved the nanopipette down to the target area at a speed of 25 μm/s and the ion current was recorded in FPGA at the same time. Based on our previous work(32), when the nanopipette gets close to the sample surface the ionic current decreases exponentially. The ionic current vs tip-surface distance curve is shown in Fig. S1. When the ionic current drops by 2%, the nanopipette approaches a position 200 nm above the sample surface. Once the nanopipette had approached this position, the piezo stopped moving and maintained the nanopipette position above the sample surface. Positive voltage or pressure pulses with different amplitudes and dwell times were applied to deliver specific amounts of Alexa-555 labelled WGA molecule onto the sample surface. We found that the magnitude of voltage can be tuned more precisely using the FPGA than using a pressure pulse controlled by mechanical pump.

### α-synuclein and amyloid*-*β aggregates deposition

The procedure of protein aggregates deposition was similar to that used for Alexa-555 labelled WGA. However, different size nanopipettes and applied voltages were used. The nanopipette which used to deliver protein aggregates has an aperture size of ∼300-400 nm and was backfilled with 8 μL negatively charged 1.4 μM α-synuclein or 70 μM amyloid*-*β aggregates in PBS. A +100mV voltage bias was applied between the electrodes inside nanopipette and the bath, generating a +2nA ionic current. When the nanopipette approached the cell surface, a negative voltage pulse (−500mV, 10s for α-synuclein) or positive pressure pulse (3kPa, 1s for amyloid*-*β) was applied to drive protein aggregates out of the nanopipette. We found in preliminary experiments that amyloid*-*β aggregates were more effectively delivered by pressure. Voltage pulses were not efficient in driving aggregates from the pipette, which is likely to be related to their charge.

### Dye and α-synuclein oligomer Injection into a live cell

The injection process used the same approach procedure as for local delivery. Once the nanopipette had approached the cell surface, the piezo moved the pipette down by an extra 5 μm in one step to penetrate the cell membrane. The penetration depth was 5 μm because the nanopipette needed enough force to penetrate the cell membrane and if the depth was less than 5 μm we found that the nanopipette would not insert into the cytoplasm. The HEK cell has a height of ∼10 μm (measured by moving the focal plane between the upper and bottom cell surface), so a penetration depth larger than 5 μm resulted in damage to the cell and potentially crushing of the nanopipette tip. Once the nanopipette had penetrated the cell membrane, a voltage pulse (−500mV, 10s) was applied to force the molecules out of the nanopipette and into the cytoplasm. After the injection process was finished, the nanopipette was retracted back by 5 μm in one step preventing the penetrated cell sticking on the nanopipette and moving up with it.

### Light sheet imaging

The detailed experimental procedure of single objective cantilever SPIM (socSPIM) imaging has been published(20). socSPIM was implemented on a standard inverted microscope (Eclipse TiU, Nikon) with an internal magnification of 1.5×. Fibre-coupled 488 nm (15 mW iFLEX2000, Qioptic), 561 nm (50 mW DPL561, Cobolt) and 640 nm (20 mW iFLEX-2000, Qioptic) diode laser lines and excitation filters (FF01488/625, LL02-561-25 and FF01-640/14-25 Semrock) were used. A cylindrical lens was used to generate the light sheet and a commercial AFM cantilever (ContAl-G, BudgetSensors) which has a reflective aluminum coating was attached onto a machined brass rod using cyanoacrylate adhesive to reflect the light sheet onto the cell sample. Either a 100× (150x with internal magnification) 1.49 numerical aperture (NA) oil immersion objective lens (MRD01991, Nikon) or a 60x (90x with internal magnification) 1.27 NA water immersion objective lens (MRD01991, Nikon) was used to focus the light sheet to the cell and collect the fluorescence emission. The emitted fluorescence was filtered by emission filters (LL02-561-25, Semrock or #67-038, Edmund Optics) and focused onto the EMCCD camera (Evolve 512 Delta, Photometrics) using a tube lens. The sample stage was controlled by a xyz piezo (P-611.3 Nanocube, Physik Instrumente) to enable 3D scanning with a scanning step of 100nm and frame rate at 0.2 Hz. The scanning data was first assembled as a hyperstacks and then reconstructed as a 3D projection or 3D volume rendering using the 3D projection function and 3D viewer plugin in FIJI where the threshold was 40 and resampling factor was 1, respectively.

### TIRF and HILO imaging

Fibre-coupled 488 nm (15 mW iFLEX2000, Qioptic), 561 nm (50 mW DPL561, Cobolt) and excitation filters (FF01488/625 and LL02-561-25 Semrock) were used. A 100× (150x with internal magnification) 1.49 numerical aperture (NA) oil immersion objective lens (MRD01991, Nikon) was used to focus the laser beam and collect the emission fluorescence. The laser beam was incident at an angle larger than the critical angle of glass/water interface when focused at the back focal plan of objective lens to achieve total internal reflection. When using HILO mode the incident angle was smaller than TIRF and an iris was used for optimal section. TIRF and HILO were used when the functions of local delivery and intracellular injection were characterized.

### Cell culture

HEK cells and MyD88-/- macrophages with GFP labeled MyD88 were gifts from Dr Lee Hopkins and Prof. Clare Bryant. They were cultivated in 1 x T175 flask *with* DMEM-Complete medium (DMEM, 10% FCS, 2 mM L-glutamine and 100 U/ml penicillin/100 mg/ml streptomycin) at 37°C/5% CO2. The lentiviral transfection process was carried out in a five-days procedure as describe before(10).

### α-synuclein and amyloid*-*β preparation

70 μM α-synuclein monomer, expressed and purified in E.coli(33–35) was shaken at 37 °C for 24 hours in order to produce a solution containing α*-*synuclein aggregates. The amyloid*-*β was purchased from American Peptide (Beta Amyloid 1-42, 1.0mg). 1 mg of amyloid*-*β pure peptide was dissolved in Hexafluoro-2-propanol and incubated at room temperature until a clear solution was formed. The solution was then aliquoted and stored at −80 °C. When required, the solution was thawed and left open in a Fume Hood overnight to evaporate the HFIP, leaving a peptide pellet. The peptide was dissolved in DMSO and then transferred to a new eppendorf with DMEM. The solution was diluted in DMEM buffer (DMEM + 1% FCS + 2 mM L-glutamine). The aggregation mixture was incubated for 6 hours at 37 °C with constant shaking of 200 r.p.m (New Brunswick Scientific Innova 43, 25 mm orbital diameter) and centrifuged at 14.2 r.p.m. for 10 min to remove any fibrillar pellets. amyloid*-*β fibrils were formed by aggregation for 60 hours. Before nanopipette delivery and injection, both α-synuclein and amyloid*-*β aggregates solution are sonicated for 20 min to ensure that there were no large aggregates that could block the nanopipette.

## Results

### Local delivery selective plane illumination microscopy

To address the difficulty of delivering precise amounts of molecules or even single molecules to a specific cell and imaging them simultaneously, we developed a local delivery selective plane illumination microscopy (ldSPIM). ldSPIM was implemented on an inverted optical microscope using a nanopipette under distance feedback control for delivery of molecules. An AFM cantilever attached to a machined brass rod to introduce a light sheet to the cell sample so that only a single objective was needed for the experiment (Fig. 1(a) and (c)). The nanopipette used for delivery approached the cell surface at an angle of 45o and was stopped at a height of approximately 200 nm above the cell surface as determined by the ionic current dropping by 2% (Fig. 1(b)). From this position molecules were delivered to the sample surface or intracellularly injected into cytoplasm driven, either by voltage or pressure. The AFM cantilever was positioned vertically next to the cell to reflect the light sheet (Fig. 1(b)). The axial thickness and Rayleigh of the sheet could be calculated by imaging the fluorescence without the cylindrical lens, as was done previously(20). The light sheet had a thickness of 1.63 μm FWHM with a Rayleigh length of 4.3 μm using a 60x 1.27NA water immersion objective. The sample stage was mounted onto a xyz piezo to enable whole-cell 3D scanning.

**Figure 1.**
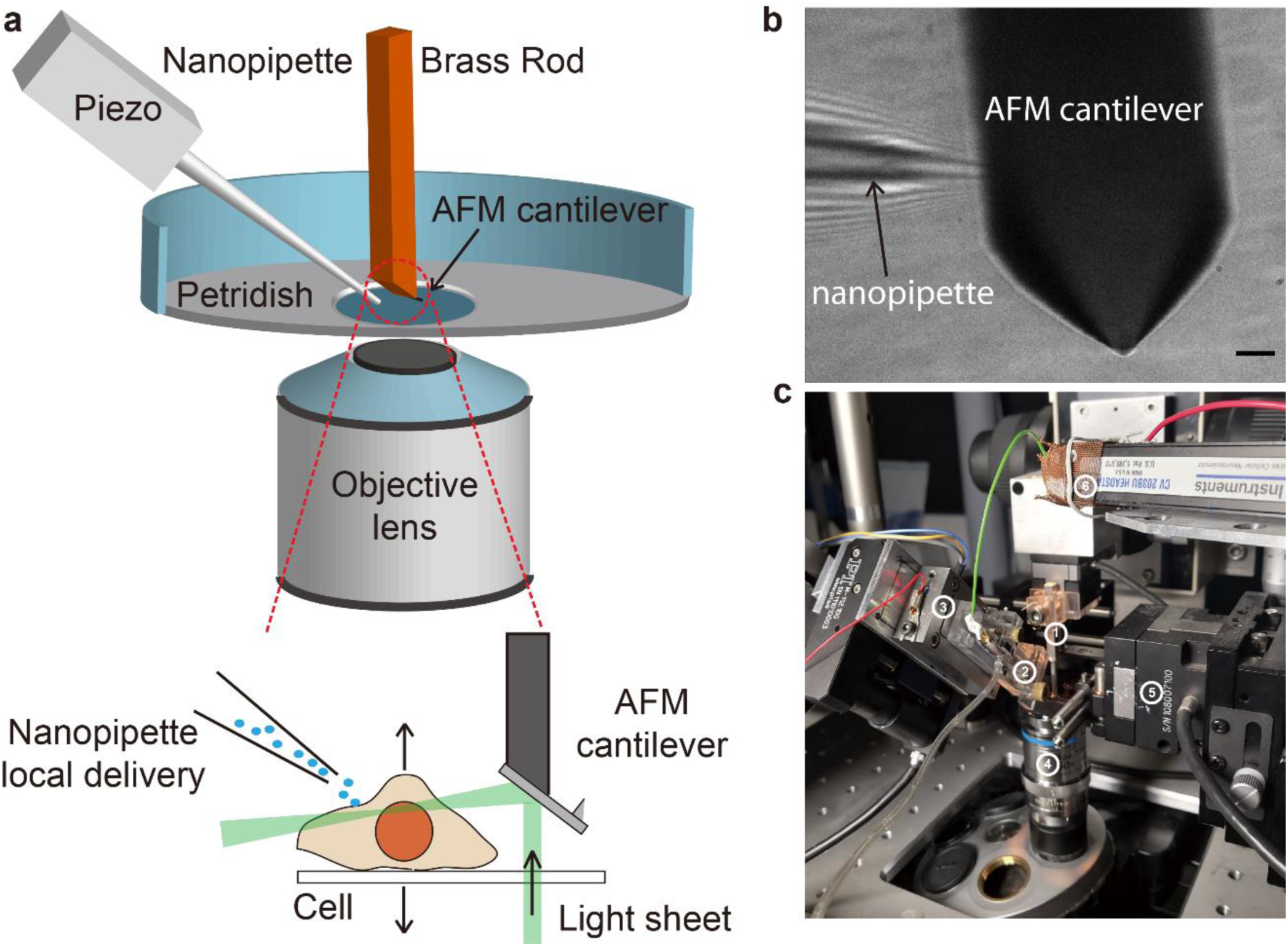
Principles of ldSPIM. **(a)** the schematic of the ldSPIM set up: Piezo controlled nanopipette and reflective AFM cantilever are combined together to locally deliver target molecules onto the sample surface and visualize the sample in a fast 3D live imaging mode. **(b)** the image of the nanopipette and the cantilever under the brightfield microscope. Scale bar is 10 μm. **(c)** Photograph of the ldSPIM setup in use. 1) is the metal rod, 2) is nanopipette on a holder, 3) is the Piezo, 4) is objective lens, 5) is the sample stage and 6) is the detecting head of pitch clamp amplifier.

We performed a series of proof-of-concept experiments to demonstrate that ldSPIM provided four fundamental capabilities: 1) ldSPIM is able to locally deliver target molecules to a pre-defined position on sample surfaces; 2) the amounts of delivered molecules can be precisely controlled. 3) ldSPIM is capable of delivering single molecules. 4) ldSPIM can penetrate the cell membrane to deliver molecules into cytoplasm.

### Local delivery to a specific position

To demonstrate the accuracy of local delivery we used the nanopipette to deposit Alexa-555 labelled WGA molecules at 24 dispersed coordinates in a circle pattern and 25 dispersed coordinates of a star pattern on a glass surface, one by one. If ldSPIM is able to deliver WGA molecules to these predefined positions, the fluorescence signal of each point should replicate the pattern. The results in Fig. 2(a) and (b), show that the nanopipette delivered the Alexa-555 labelled WGA molecules to form the circle and star patterns (the full video is in Movie S1 and S2). Furthermore, the actual deposited areas overlap well with the predefined coordinates labelled in blue spots, indicating that the nanopipette precisely delivered WGA molecules to a specific position.

**Figure 2.**
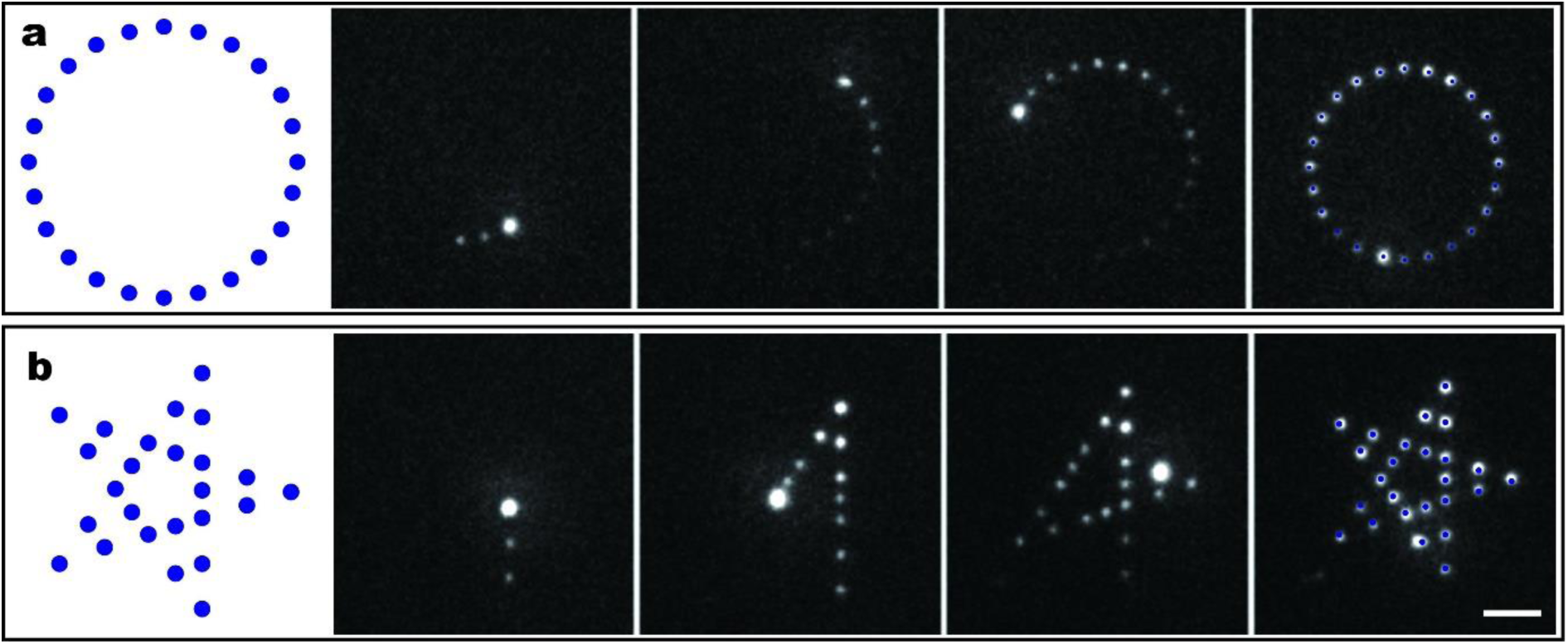
Accurate delivery of Alexa-555 labelled WGA molecules to specific position on glass surface forming different patterns. The coordinates of a circle and star, the delivering process and the overlay of coordinates and real deposited WGA molecules are shown in **(a)** and **(b)**. The scale bar is 20 μm.

### Controlling the amount of molecules deposited

When a voltage pulse is used to deposit molecules onto a sample surface, there are three parameters that can affect the amount of delivery: the concentration of Alexa-555 labelled WGA molecules in the PBS buffer, the magnitude of the voltage pulse and the pulse dwell time. When a nanopipette is loaded with a certain concentration of WGA solution, it cannot easily be washed and reloaded with a different concentration of WGA solution. Therefore, if we want to use a different concentration of Alexa-555 labelled WGA PBS buffer, we must keep replacing the nanopipette. However, this introduces the inner diameter of the nanopipette as another variable. Based on these two facts, the concentration was kept constant and the magnitude of voltage pulse and the pulse dwell time independently varied in experiment on fixed HEK cell surface depositing Alexa-555 labelled WGA molecules. The dwell time was first set at 2s and the magnitude of voltage was varied from 0mV to 900mV. The fluorescence of deposited Alexa-555 labelled WGA molecules on fixed HEK cell surface increased linearly with the increase in the voltage pulse (Fig. 3(a)). The effect of changing the pulse time from 0s to 7s was then quantified, with the pulse voltage set at a constant 600mV. The fluorescence of deposited molecules also showed a linear relationship with dwell time (Fig. 3(b)). These results demonstrate that the amount of deposited Alexa-555 labelled WGA molecules can be precisely controlled by varying the voltage or dwell time.

**Figure 3.**
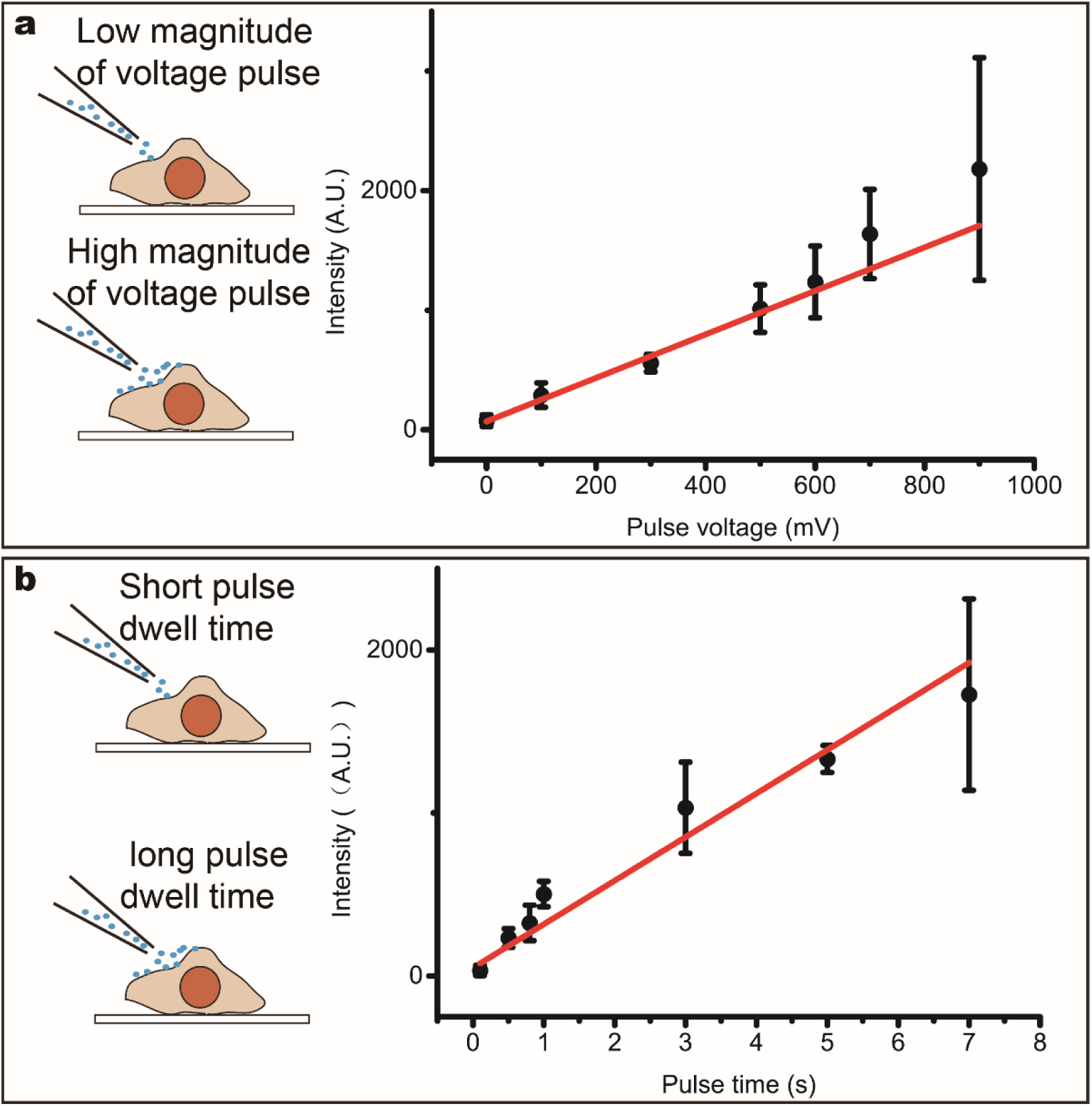
The amount of Alexa-555 WGA molecule deposited on fixed HEK cell surface is fine-tuned by adjusting the magnitude of voltage pulse and pulse dwell time. Here we use the fluorescence of Alexa-555 WGA molecules deposited on fixed HEK cell surface to represent the amount. **(a)** When the pulse dwell time is fixed at 2s, the fluorescence of deposited Alexa-555 WGA molecule increases linearly with the raising of pulse voltage from 0 mV to 900mV. **(b)** When the pulse voltage is a constant of 600mV, the fluorescence of deposited Alexa-555 WGA molecule also has a linear relationship with the increase of pulse dwell time from 0s to 7s. The error bar in both **(a)** and **(b)** represents the standard deviation of 6 repeats of each parameter on the same fixed HEK cell.

### Single-molecule delivery

The capability to deposit single molecules onto the cell surface was demonstrated by depositing Alexa-555 labelled WGA molecules onto the surface of fixed HEK cell. Using a set of delivery parameters for single-molecule delivery, a single spot was delivered onto a fixed HEK cell surface and then was imaged until it photobleached (Fig. 4(a) and supplementary Movie S3). The number of molecules delivered was determined by observing the photobleaching of Alexa-555 (Fig. 4(b)). In Fig. 4b, two photobleaching steps could be observed, indicating that there are two Alexa-555 fluorophores on the WGA molecules. Since the degree of labelling of the WGA was 1.6 on average, this result demonstrates the ability to deliver single WGA molecules. The single-molecule deposition protocol was repeated 30 times using the auto delivery and imaging algorithm and over half of the replicates showed that on average only one or two single Alexa-555 labelled WGA molecules were deposited on the fixed HEK cell surface (Fig. 4(c)).

**Figure 4.**
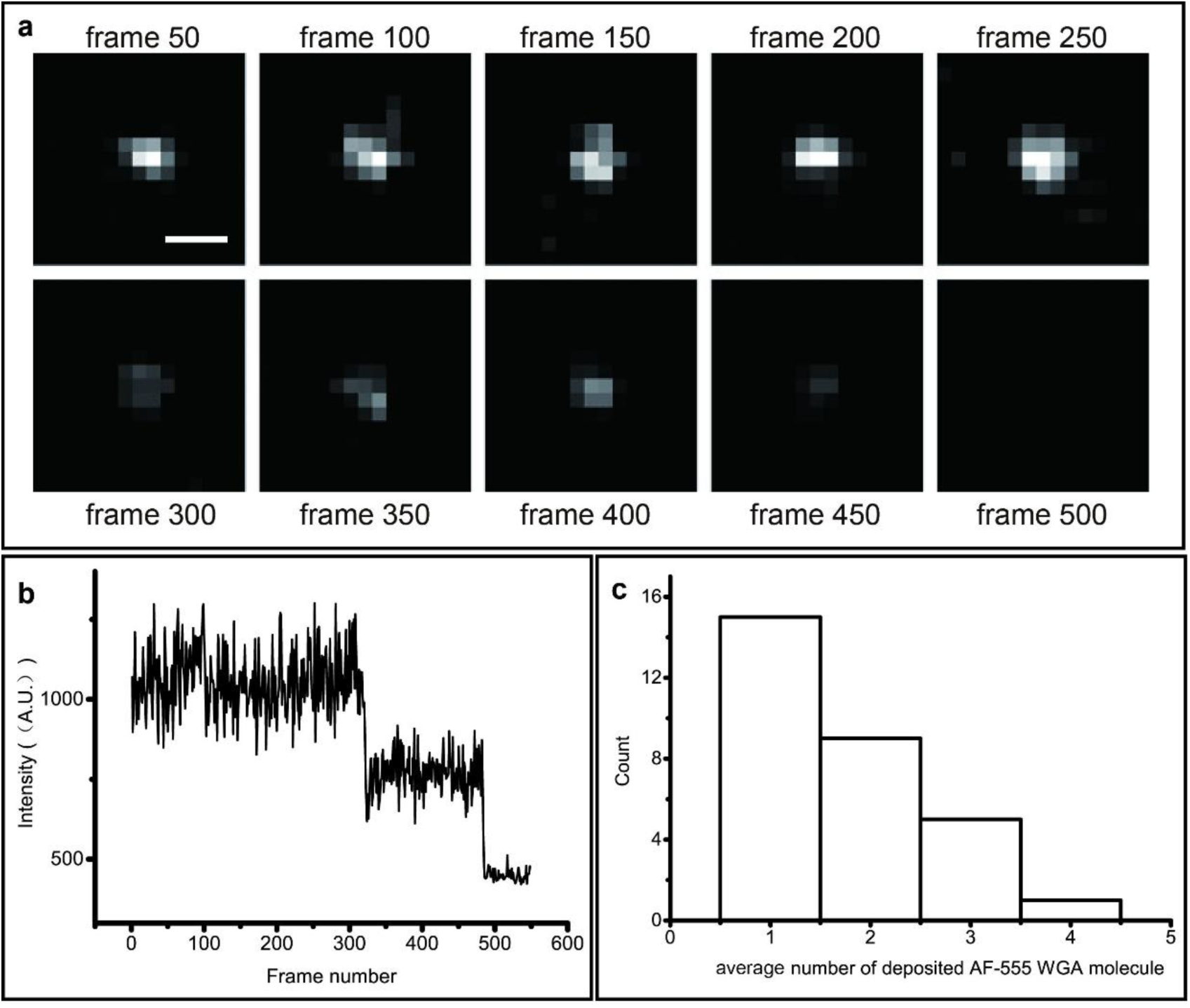
Single Alexa-555 labelled WGA molecules deposited on fixed HEK cell surface. **(a)** Photobleaching of a deposited WGA molecule. **(b)** Intensity profile of the WGA molecule bleached in **(a)**. The intensity decreases in two steps suggesting the presence of two Alexa-555 dye molecules. As the degree of labelling of our WGA molecules is 1.6:1, this approximately corresponds to a single WGA molecule in **(a). (c)** The distribution of the average number of deposited Alexa-555 WGA molecule over thirty replicates. The scale bar is 500 nm.

### Intracellular injection

Besides surface delivery, ldSPIM is also able to achieve intracellular deposition by penetrating the cell membrane, followed by direct delivery of molecules into the cytoplasm using a voltage or pressure pulse. We used Alexa-647 dye (Fig. 5(a)) and Alexa-594 labelled α-synuclein aggregates (Fig. 5(b)) to demonstrate that ldSPIM can not only introduce small molecules but also relevant big protein oligomers or aggregates inside the cytoplasm. Before nanopipette penetration, the cell was black without any fluorescence. Once the nanopipette was inserted into the cell, the fluorescence at the tip of nanopipette could be identified as a bright spot inside the cell. During the injection process, the cell was illuminated with fluorescence from Alexa-647 and Alexa-594 labelled α-synuclein aggregates, while the background remanded dark. The fluorescence intensity profile of the injected cells changed synchronously with the injection process (Fig. 5(c) and (d)). Alexa-647 is small molecule so the fluorescence was uniform and was observed over the whole cell including the nucleus. In contrast, the larger synuclein aggregates could not enter the nucleus. There are also some brighter spots left over the injection. Since the size of nanopipette’s aperture is around 100 nm, it can specifically introduce large aggregates into cytoplasm without affecting the nucleus. The videos of injection are shown in supplementary Movie S4 and S5.

**Figure 5.**
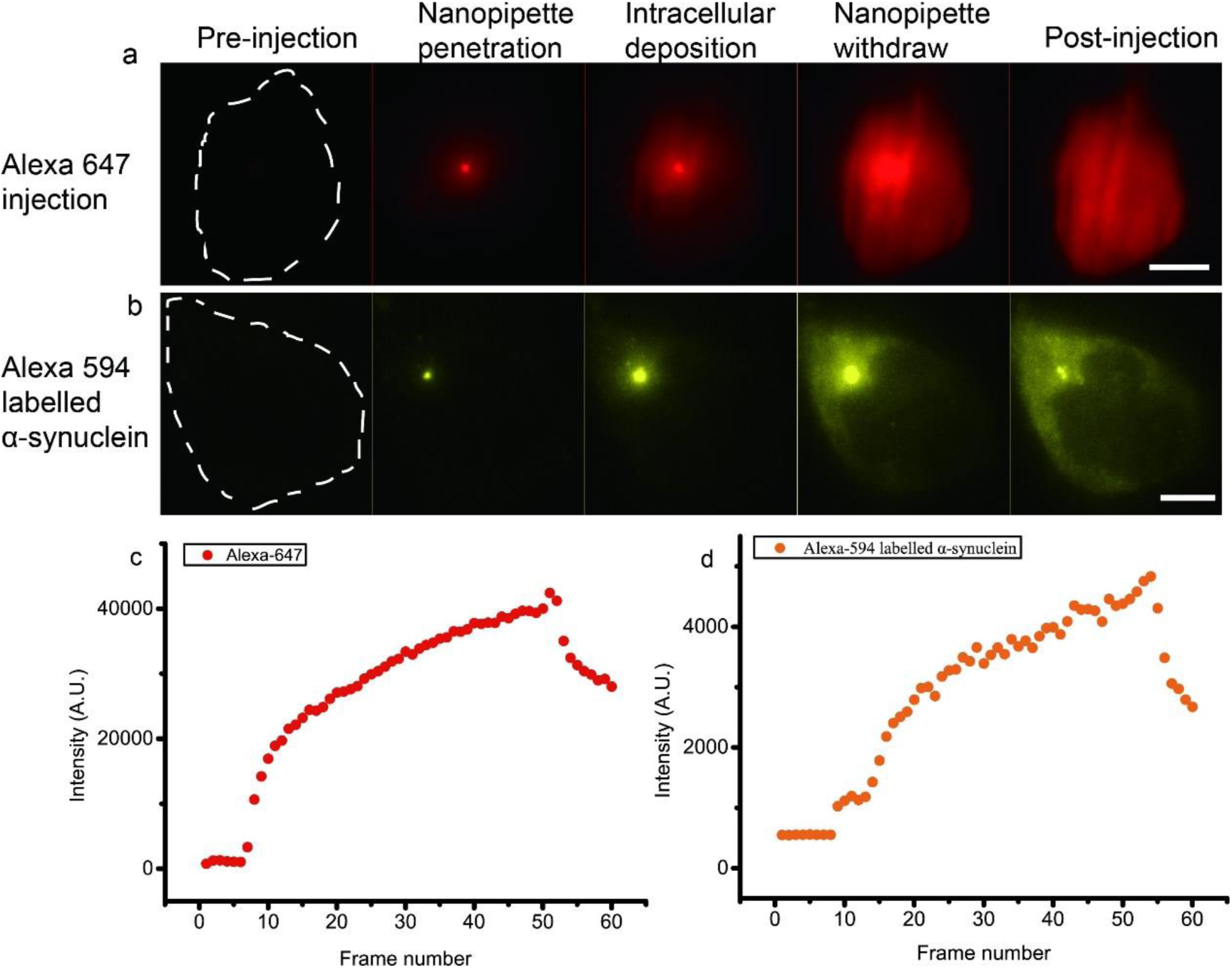
Nanopipette delivery into live cells. Intracellular deposition of Alexa-647 and Alexa-594 labelled α-synuclein aggregates over time are shown in **(a)** and **(b)**. The dashed lines indicate the outline of the cells. The fluorescence intensity of the deposited molecules inside the cell over injection process are shown in **(c)** and **(d)**. The fluorescence intensity increased near frame 10 and kept rising until frame 50 representing the nanopipette penetration and continuous deposition of Alexa-647 and Alexa-594 labelled α-synuclein aggregates. The decrease near frame 55 indicates the withdrawing of nanopipette. The scale bar is 4 μm.

### 3D imaging of Myddosome formation

The activation of innate immune system and the subsequent downstream responses, such as inflammation, also play key roles in many disease progressions. Preclinical and clinical studies have shown that the activation of the innate immune system could be involved in the pathologies of neurodegenerative diseases(36–40). In Alzheimer’s disease for instance, both small soluble protein aggregates assembled from amyloid-β (oligomers) and larger fibrillary aggregates are able to bind TLR4 and other receptors on microglia, the resident macrophage within the brain, to activate the immune system and trigger downstream inflammation responses. According to our recent research, the TLR4-mediated inflammatory triggered by amyloid*-*β aggregates could be a major pathophysiological pathway in AD(41).

The core concept of ldSPIM is that it can deposit molecule to a target cell while the light-sheet illumination allows the cell response to be visualized at single-molecule level in 2D or 3D mode. Therefore, we used ldSPIM to deposit amyloid*-*β aggregates on the cell surface to trigger macrophages expressing a MyDD88-eGFP fusion construct and we visualized the formation of Myddosome in live cells in 3D. In this case, we used pressure to deliver amyloid*-*β aggregates onto macrophages. Compared with amyloid*-*β monomer triggered macrophages that showed no response (Fig. 6(a) and (b), Supplementary Movie S6 and S7), the macrophage triggered by amyloid*-*β aggregates formed Myddosome underneath the plasma membrane within 300 seconds (Fig. 6(c) and (d) Supplementary Movie S8 and S9). The 3D light sheet scanning also enables a 3D rendering of formed Myddosome to show the spatial distribution (Fig. 6 (e) and Supplementary Movie S10). Moreover, the complete Myddosome action triggered by amyloid*-*β aggregates from assembling, splitting and then disassembling could be observed in Fig. 6(f) and (g).

**Figure 6.**
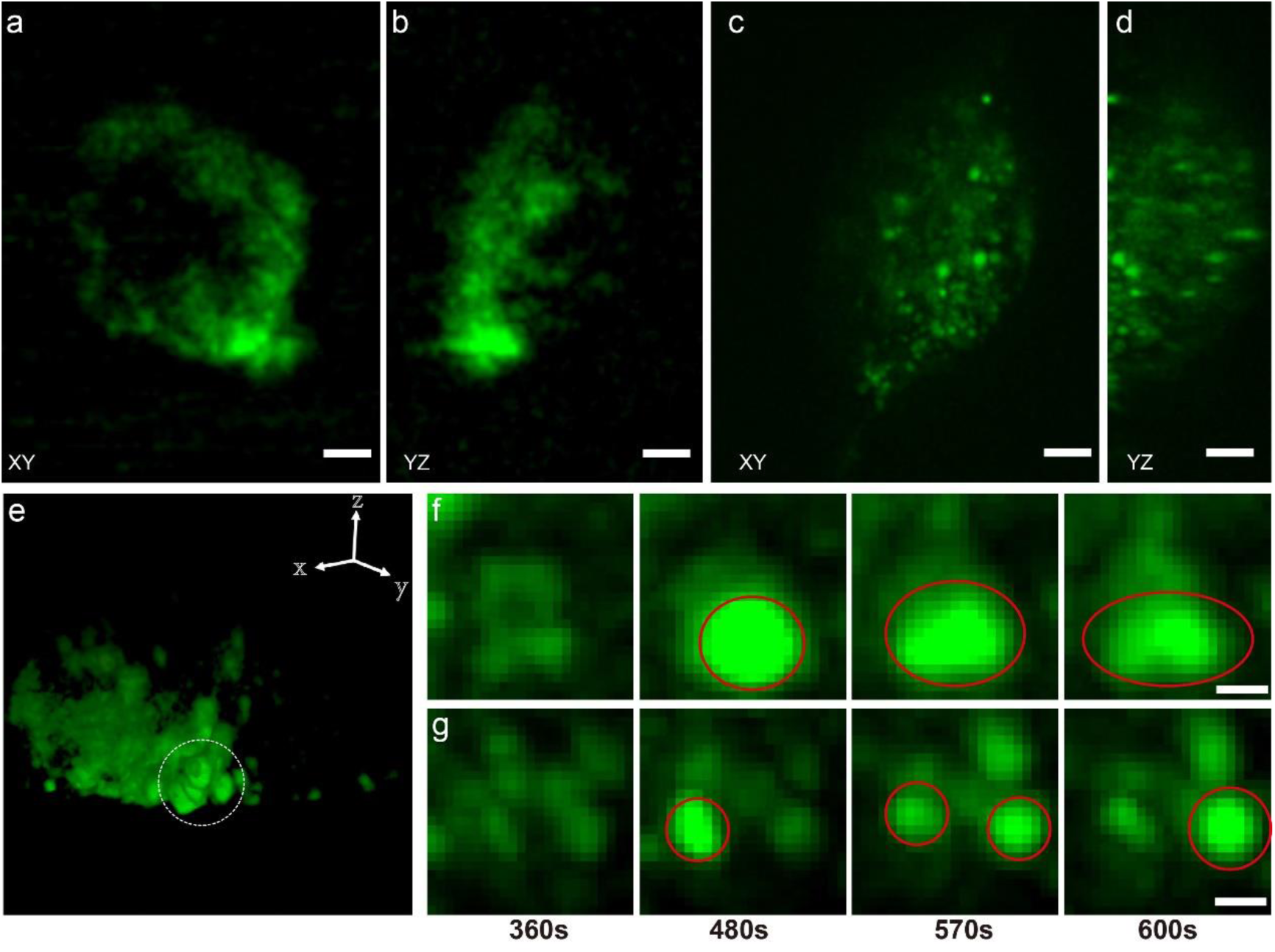
Representative images of whole-cell 3D imaging of Myddosome formation in live macrophages (n=10). **(a) and (b)** the intensity projection of amyloid*-*β monomer control in XY and YZ at t=300 s. **(c) and (d)** the intensity projection of Myddosome formed in macrophage triggered by amyloid*-*β aggregates in XY and YZ at t=300 s. The scale bar is 4μm. **(e)** the 3D volume rendering of the data in **(c)** showing the spatial distribution of Myddosomes. Time course of selected Myddosome in **(e)** response to amyloid*-*β triggering are shown in **(f)** and **(g). (f)** the process of Myddosome forming and splitting. **(g)** the process of Myddosome assembling and disassembling. 480 s post amyloid*-*β aggregates activation, Myddosome with different sizes formed. Then at 570s post triggering, the big Myddosome (in **(f)**) started splitting while the small one (in **(g)**) began to disassemble. At 600 s post triggering, the big Myddosome was still splitting and the small one had disassembled. The scale bar is 2 μm.

## Discussion

We have demonstrated that the ldSPIM is capable of delivering single or precise amounts of biomolecules to specific locations on a single live cell. The experiments also show that ldSPIM can perform intracellular injection of dye molecules as well as protein aggregates. The most notable advantage of ldSPIM is that it is the first instrument being able to perform precise local delivery in combination with high-resolution light-sheet microscopy with single molecule sensitivity. Myddosomes formation induced by TLRs triggering has been imaged recently(10). However, this was done using TIRF. Using ldSPIM, the Myddosome formation triggered by amyloid*-*β aggregates was visualized in live cells with real-time whole cell 3D imaging. The capability to precisely deliver molecules onto the apical surface of a single cell means that it is now possible to study the early events of immune cell triggering in a controllable and reproducible way. The instrument also opens up a wide range of applications for studying cell signaling using nanopipette delivered agonists but also delivering labelled molecules into the cell and then following where they go and how the cell responds. For example, we can deliver labelled protein aggregates into the cell and determine if they bind to organelles or are removed by the proteasome or autophagy. Overall, this instrumentation should open up a range of new single-molecule experiments on live cells.

In the future the ldSPIM can be improved in two additional aspects. Firstly, the ldSPIM uses a single barrel nanopipette, which is only able to deliver one kind of molecules to a location at a time. If two reagents are required to be delivered to a same position, two different nanopipette must be used to finish the deposition separately. This will make time sensitive experiments very difficult and introduce the pipette size as an extra variable. Secondly, the light sheet used in ldSPIM is reflected by an AFM cantilever which occupied the vertical space above the sample. It is challenging to add another detection method such as patch-clamp due to the limited space. A double barrel nanopipette(42) will be used to address the first issue. The double barrel nanopipette has two independent channels in one capillary so it is able to deliver two kinds of molecules e.g. AF594-labelled α-synuclein and Alexa-647 labelled amyloid*-*β to one or more locations e.g. cytoplasm and nucleus at the same time. Optimizing the optical pathway to achieve a new version of light sheet microscopy(43) without using a mirror to reflect the beam and removing the AFM cantilever will release more space, allowing more functionality in a single experiment and compatibility with well plates for high-throughput studies.

## Author Contributions

B.Li, A.Ponjavic, W.Chen and D.Klenerman designed the research. B.Li implemented the nanopipette delivery and injection devices. A.Ponjavic implemented the single objective cantilever light sheet microscope. B.Li performed the local delivery and injection and Myddosome formation experiments. L.Hopkins, C.Hughes, Y.Ye and C.Bryant prepared all the cells needed in each experiment. B.Li, A.Ponjavic and D.Klenerman wrote the manuscript with input from all authors.

## Acknowledgement

This work was supported by the UK Dementia Research Institute which receives its funding from DRI Ltd, funded by the UK Medical Research Council, Alzheimer’s Society and Alzheimer’s Research UK and by the Royal Society and the EPSRC (EP/L027631/1).

